# Evaluation of ready-to-use freezer stocks of a synthetic microbial community for maize root colonization

**DOI:** 10.1101/2023.05.10.540175

**Authors:** J. Jacob Parnell, Simina Vintila, Clara Tang, Maggie R. Wagner, Manuel Kleiner

**Affiliations:** Department of Plant and Microbial Biology, North Carolina State University, Raleigh, NC, United States; Department of Ecology and Evolutionary Biology, University of Kansas, Lawrence, KS, United States; Kansas Biological Survey & Center for Ecological Research, University of Kansas, Lawrence, KS, United States

**Author notes:** These authors contributed equally to this work (J. Jacob Parnell, Simina Vintila).

**Keywords:** Synthetic communities, maize, root colonization, plant-microbe interactions, plant microbiome, SynCom

## Abstract

Synthetic microbial communities (SynComs) are a valuable tool to study community assembly patterns, host-microbe interactions, and microbe-microbe interactions in a fully controllable setting. Constructing the SynCom inocula for plant-microbe experiments can be time consuming and difficult because a large number of isolates with different media requirements and growth rates are grown in parallel and mixed to appropriate titers. A potential workaround to assembling fresh SynCom inocula for every experiment could be to pre-make and freeze SynComs on a large scale, creating ready-to-use stock inocula. The objective of this study was to compare the reproducibility, stability, and colonization ability of freshly prepared versus frozen SynCom inocula. We used a community of seven species known to colonize maize roots. The results from inoculation with the frozen SynCom were as consistent as standardized *de novo* construction of fresh SynCom. Our results indicate that creating frozen SynCom inocula for repeated use in experiments not only saves time, but could also improve cross-experiment reproducibility. Although this approach was only validated with one SynCom, it demonstrates a principle that can be tested for improving approaches in constructing other SynComs.

**Importance:** Synthetic communities (SynComs) are an invaluable tool to characterize and model plant-microbe interactions. Multimember SynComs approximate intricate real-world interactions between plants and their microbiome, but the complexity and time required for their construction increases enormously for each additional member added to the SynCom. Therefore, researchers who study a diversity of microbiomes using SynComs are looking for ways to simplify the use of SynComs. In this manuscript, we evaluate the feasibility of creating ready-to-use freezer stocks of a well-studied seven-member SynCom for maize roots. The frozen ready-to-use SynCom stocks work according to the principle of “just add buffer and apply to sterilized seeds or seedlings” and thus can save multiple days of laborious growing and combining of multiple microorganisms. We show that ready-to-use SynCom stocks provide comparable results to freshly constructued SynComs and thus allow for large time savings when working with SynComs.

## Introduction

Plant-associated microbiota play key roles in plant evolution, development, health, and stress resilience; studying these roles is critical for understanding fundamental principles of plant biology and developing and applying biotechnology for sustainable agriculture (Compant et al., 2019; Mendes et al., 2011; Vandenkoornhuyse et al., 2015). Microorganisms improve plant host fitness by facilitating nutrient uptake (Zhang et al., 2021), providing defense against pathogens (Carrión et al., 2019; Durán et al., 2018), and alleviating abiotic stress (Berendsen et al., 2012). Dissecting and using the microbial mechanisms that benefit plants is becoming more important as rising global population and climate change place greater demands on sustainable agricultural production (Parnell et al., 2016). Although the initial assembly of plant-associated microbiota is influenced by stochastic processes, mounting evidence suggests that healthy plants can select particular microorganisms to establish beneficial communities that are complex and structured, yet reproducible (Finkel et al., 2017; Knief et al., 2010; Müller et al., 2016; Walters et al., 2018).

Synthetic microbial communities (SynComs) that reduce the complexity of the plant-associated microbiota while maintaining key structures and functions allow for hypothesis-driven experiments with reproducible conditions (Liu et al., 2019; Müller et al., 2016; Vorholt et al., 2017). Experiments using SynComs in gnotobiotic plant systems have described assembly patterns resulting from specific plant-microbe (Bodenhausen et al., 2014) and microbe-microbe interactions (Snelders et al., 2020, Coker et al., 2022), identified keystone species and assembly patterns (Carlström et al., 2019; Niu et al., 2017), determined microbial niche specialization (Bai et al., 2015) and led to the discovery of microbe-dependent heterosis in maize (Wagner et al., 2021). In short, SynComs are a powerful tool for unraveling complex plant-microbe and microbe-microbe interactions and defining the plant holobiont.

Reproducible construction of even mildly complex SynComs requires significant time and labor. Recent evidence highlights the importance of reproducible SynCom construction, suggesting that the inoculation ratio of community members influences microbial community interactions (Aharonovich and Sher, 2016; Gao et al., 2021; Marín et al., 2021; Venturelli et al., 2018). Additionally, time- and labor-intensive construction of SynComs would become cost-prohibitive for SynComs developed as commercial products for widespread application in agriculture. In order to improve consistency between experiments, and extend concepts and plant-beneficial mechanisms to agricultural production, new approaches to building SynComs are required that are less labor intensive while providing consistent ratios of SynCom members.

Here we explore the stability and efficacy of fresh and frozen aliquots of a seven member SynCom that has been developed as a simplified and representative SynCom for maize (Niu et al., 2017). We used root colonization following inoculation on maize seeds to measure stability and efficacy.

## Materials and Methods

### Preparation of fresh and frozen SynCom inocula

The synthetic community used in this study contained the following bacterial species: *Stenotrophomonas maltophilia* AA1 (ZK5342), *Brucella pituitosa* AA2 (ZK5343), *Curtobacterium pusillum* AA3 (ZK5344), *Enterobacter ludwigii* AA4 (ZK5345), *Chryseobacterium indologenes* AA5 (ZK5346), *Herbaspirillum robiniae* AA6 (ZK5347), and *Pseudomonas putida* AA7 (ZK5348) as described in Niu et al. (2017; see also Niu & Kolter 2018). Following those same studies, we used the “Sugar Bun” (Johnny’s Seeds, Cat. 267T) variety of maize as plant host throughout this study.

To determine the variability between different SynCom preparation events and the impact of freezing on the viability of the SynCom, we conducted two experiments in which four different SynCom mastermixes were prepared and inoculated onto plants either directly (fresh) or after freezing. To determine if the duration of freezing had an impact on the SynCom viability, we tested the mixes after one hour and after one week of freezing (Figure 1). We constructed the synthetic community following the previously published protocol by Niu and Kolter (2018) with some modifications. We streaked freezer stocks of each species onto selective 0.1x tryptic soy agar plates with species-specific antibiotics and then incubated the plates at 30°C for two days. Individual colonies from plates were inoculated into 5 ml of tryptic soy broth without dextrose (VWR) and shaken for 8 hours at 30°C. We transferred 0.1 ml of each species’ culture to a separate 250 ml flask with 125 ml of tryptic soy broth and flasks were shaken at 30°C for 14 to 16 hours. We centrifuged the cultures at 8,000 x g for 10 min at 4°C. Cell pellets were washed with 5 ml of phosphate buffered saline (PBS) pH 7.4 (Fisher Scientific), re-pelleted by centrifugation at 8,000 x g for 10 min at 4°C, and re-suspended in 10 ml PBS. We combined species to build a SynCom master mix with *S. maltophilia* and *P. putida* at 10^7^ cells/ml and all other species at 10^8^ cells/ml. Concentrations were estimated from optical density (OD) readings calibrated to cells/ml using direct cell counting under a microscope.

**Figure 1.**
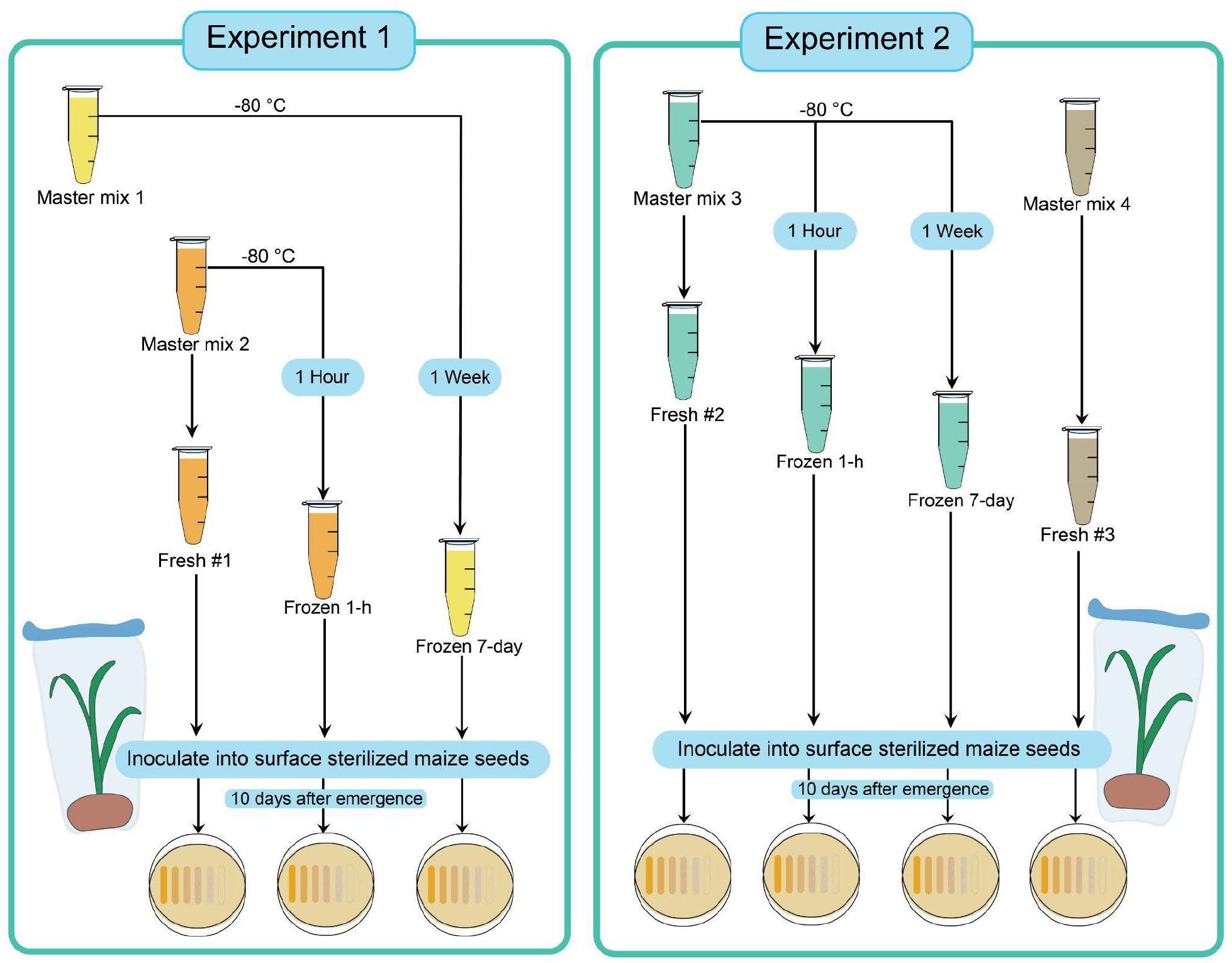
Experimental set up of master mixes and their use in Experiment 1 and 2.

The SynCom master mix was divided into aliquots for plate counting, creating freezer stocks, and direct inoculation of the fresh community onto sterilized maize seeds. We prepared fresh and frozen, ready-to-use stocks of the SynCom in the same exact way as follows: 1 ml of the master mix was added to cryotubes containing 0.7 ml of sterilized PBS:glycerol (1:1) yielding a final volume of 1.7 ml with 20% glycerol. The fresh stocks were never frozen. To use a stock, 1.7 ml of the stock was mixed with 8.3 ml PBS to yield a 10 ml inoculum with a final concentration of 10^6^-10^7^ cells/ml. The resulting 10 ml of 10^6^-10^7^ cells/ml SynCom mix (fresh or frozen) was added per liter of 0.5x Murashige & Skoog medium (Caisson labs) yielding a final concentration of 10^4^-10^5^ cells/ml of each bacterial species. This mix was then used for plant inoculation as described below. Four replicate SynCom master mixes were created for two experiments over the course of the study.

In the first experiment (Experiment#1), a SynCom mix (Master mix 1) was prepared and frozen at -80 °C. One week later, another SynCom mix (Master mix 2) was prepared and split into a fresh and a frozen aliquot. The fresh aliquot from Master mix 2 was immediately inoculated into surface sterilized maize seeds (Fresh #1). At the same time, the frozen aliquot from Master mix 1 was thawed and inoculated (Experiment 1: Frozen 7-day). The frozen aliquot of Master mix 2 was frozen at -80 °C for one hour, thereafter the mix was thawed and inoculated on the same day (Experiment 1: Frozen 1-h). In the second experiment (Experiment#2), two more SynCom mixes were prepared separately (Master mix 3 and Master mix 4). Master mix 3 was split into three aliquots one to be inoculated fresh the same day (Fresh #2), after being frozen for one hour (Experiment 2: Frozen 1-h), and after being frozen for one week (Experiment 2: Frozen 7-day). Master mix 4 was prepared and inoculated fresh (Fresh #3) on the day that Frozen 7-day (Experiment 2) was inoculated into surface sterilized maize seeds.

### Inoculation of surface sterilized of maize seeds and growth

In a laminar flow hood, we surface sterilized Sugar Bun seeds by submerging them in 70% (v/v) ethanol for 3 minutes, followed by submerging them in 2% (v/v) sodium hypochlorite solution for 3 minutes, and then washing them 10 times with sterilized deionized water. The final wash was plated onto 0.1X TSA plates to test for residual contamination after the sterilization procedure as described by Niu et al (2017). We inoculated 15 surface sterilized seeds with the fresh and 15 surface sterilized seeds with the 1 hour frozen SynCom inocula as previously described (Wagner et al., 2021). Surface sterilized seeds were placed in sterile 7.5” x 15” Whirl-pak bags (Nabisco) filled with 150 ml of a calcined clay (“Pro’s Choice Rapid Dry”; Oil-Dri Corporation) and inoculated with 90 ml of 0.5x Murashige & Skoog medium containing the SynCom. We sealed the bags with sterile Aerafilm breathable film (Excel Scientific, Inc.) to keep the system sterile while allowing for aeration. Bags were randomized and placed on a shelf with light-emitting diode (LED) growth lights (16 h days, 23 °C, ambient humidity). Emergence of seedlings was documented daily and plants were harvested at 10 days post-emergence.

### Root harvest and absolute quantification of live microbial cells colonizing the roots

Roots from 9-10 plants were harvested for the colonization assay according to Niu and Kolter (2018) with some modifications. Plants were removed from bags and the roots were gently rinsed in deionized water. The whole primary root from each plant was harvested and fresh weights were recorded (200-500 mg fresh weight). Roots were cut into small pieces with a sterile knife and vortexed in a 2 ml tube for 3 minutes in 1 ml of sterile PBS with six 3 mm sterilized glass beads to recover microbial cells from the root surface as described in Niu and Kolter (2018). We serially diluted the cell suspension in PBS (10^−1^-10^−8^) and plated the dilutions on species-specific selective media as previously described (Niu and Kolter 2018). After incubation for the required time for selective growth (16-60 h at 30°C), colonies were counted and counts normalized against root fresh weight.

### Viability of SynCom members following longer-term freezing

To test the impact of freezing on the SynCom mix, 4 vials of ready-to-use stocks, which had been frozen for 60 days, were each diluted to 10^6^-10^7^ cells/ml in PBS and a dilution series (10^−1^-10^−8^) was prepared for each mix. 10 ul of each dilution in the series was plated in technical triplicates on selective plates according to Niu et. al (2018). Plates were incubated for 16-60 h at 30°C and the resulting colonies were counted.

### Statistical analysis

To detect statistical differences between different SynCom master mixes, one-way ANOVA followed by Tukey HSD was performed on the dataset using RStudio (1.4.1106) statistical software (R Core Team, 2021; RStudio Team, 2020). Adjustment of p-values was done using the Benjamini-Hochberg method. For each experiment, log CFUs / g fresh weight of each species were tested separately against the type of master mix used to inoculate the plants. Assumptions of normality were satisfied by looking at the data distribution along normal Q-Q plots. Assumptions of equal variance were tested using the Bartlett test.

## Results

Four replicate SynCom master mixes were created at separate times to test for differences in maize root colonization in terms of bacterial species colony-forming unit counts due to (1) duration of freezer storage or (2) SynCom preparation. Master mixes 1 and 2 had aliquots frozen for 1 hour or 7 days. Master mix 3 was used in its fresh form, frozen for 1 hour and frozen for 7 days. Master mix 4 was used only in its fresh form as a comparison inoculation to the 7 day frozen Master mix 3 (Figure 1). Master mixes contained 10^7^ cells/ml of *Stenotrophomonas maltophilia* and *Pseudomonas putida*, and 10^8^ cells/ml of *Brucella pituitosa, Curtobacterium pusillum, Enterobacter ludwigii, Chryseobacterium indologenes*, and *Herbaspirillum robiniae*.

### Variation of species abundances in roots from plants inoculated with fresh SynComs

We inoculated sterilized seeds in calcined clay with Murashige & Skoog medium containing the SynCom bacteria and grew the plants under controlled conditions for 10 days post-emergence before we harvested roots for bacterial colony counts (see Materials and Methods). Comparing roots of plants inoculated with the same SynCom inoculum (e.g. Master Mix #1 frozen for 1 hour) showed that, for all inocula, the abundance (log CFUs/g of root fresh weight) of each SynCom species was highly variable between replicate plants. The standard deviations for species abundances in replicate plants ranged from a log-value of 0.18 (*E. ludwigii* and *C. pusillum*) to 2.93 (*H. robiniae*) (Table 1).

**Table 1.** CFUs for each species per gram of fresh maize roots harvested 10 days post-emergence (mean log-transformed values are shown; standard deviations are in parentheses). For each SynCom and condition (fresh or frozen) CFUs/g were determined for 8 to 11 plants.

|  | Experiment #1 |  |  | Experiment #2 |  |  |  |
| --- | --- | --- | --- | --- | --- | --- | --- |
|  | Master Mix #2 |  | Master Mix #1 | Master mix #3 |  |  | Master mix #4 |
|  | Fresh #1 | Frozen 1-h | Frozen 7-d | Fresh #2 | Frozen 1-h | Frozen 7-d | Fresh #3 |
| *S. maltophilia* | 1.86 (1.20) | 2.66 (1.79) | 2.82 (1.28) | 3.40 (1.76) | 2.77 (1.53) | 0.37 (1.12) | 1.34 (1.97) |
| *B. pituitosa* | 5.68 (0.38) | 5.75 (1.00) | 5.75 (1.26) | 7.93 (0.70) | 7.41 (0.79) | 5.61 (0.64) | 5.79 (0.86) |
| *C. pusillum* | 7.71 (0.32) | 7.86 (0.27) | 8.09 (0.18) | 9.04 (0.29) | 9.22 (0.34) | 7.82 (0.33) | 7.90 (0.33) |
| *E. ludwigii* | 7.18 (0.18) | 7.69 (0.83) | 7.41 (0.50) | 9.57 (1.22) | 8.56 (0.73) | 7.53 (0.35) | 7.72 (0.51) |
| *C. indologenes* | 4.09 (1.68) | 5.68 (0.71) | 4.20 (2.32) | 6.31 (1.02) | 6.63 (1.20) | 4.75 (0.78) | 4.97 (0.84) |
| *H. robiniae* | 3.93 (2.93) | 4.33 (2.77) | 5.74 (1.80) | 7.33 (0.85) | 6.20 (0.99) | 5.17 (0.49) | 4.56 (2.08) |
| *P. putida* | 0.49 (1.09) | 2.25 (1.60) | 1.55 (1.48) | 4.48 (0.92) | 3.79 (2.05) | 1.97 (1.94) | 1.19 (1.80) |

Comparing roots of plants inoculated with different master mixes showed that abundances varied considerably for some species (Tables 1 & 2). All of the species had significantly higher abundances (CFUs/g of fresh root weight) in fresh mix 2 as compared to fresh mix 3 (p < 0.05, Table 2). When comparing fresh mixes 1 and 2 again all species had significantly higher abundances (CFUs/g of fresh root weight) in fresh mix 2 as compared to fresh mix 1 (p < 0.05, Table 2) with the exception of *S. maltophilia* (Tables 1 & 2). None of the SynCom members were significantly different in terms of CFUs/g of fresh root weight between fresh mixes 1 and 3.

**Table 2.** Statistical comparison of species abundances between plants inoculated with the three fresh SynCom master mixes. Shown are adjusted p-values from ANOVA followed by Tukey HSD comparing CFUs/g fresh weight on maize roots collected 10 days post-emergence (Table 1). Significantly different comparisons (p<0.05) are marked with an asterisk (*).

|  | SynCom Member |  |  |  |  |  |  |
| --- | --- | --- | --- | --- | --- | --- | --- |
|  | *S. maltophilia* | *B. pituitosa* | *C. pusillum* | *E. ludwigii* | *C. indologenes* | *H. robiniae* | *P. putida* |
| Fresh 1 v 2 | 0.21 | 0.0000002* | <1 e-7* | <1 e-7* | 0.0016* | 0.0008* | 0.0004* |
| Fresh 1 v 3 | 0.81 | 0.93 | 0.38 | 0.21 | 0.46 | 0.27 | 0.19 |
| Fresh 2 v 3 | 0.04* | 0.0000004* | <1 e-7* | 0.000014* | 0.028* | 0.03* | 0.005* |

### Impact of freezing SynCom stocks on species abundances in the inoculum and in roots

Abundances of each SynCom species from plants inoculated with the frozen SynCom stocks compared with the corresponding fresh SynCom master mixes suggests that freezing had little impact on each species’ ability to colonize plant roots. Results from tests with two replicate master mix communities showed that only *C. pusillum* colonization was consistently altered by freezing in both replicate experiments (Table 3). For Master Mix 3, we found a significant difference in the colonization of all 7 species when comparing the fresh (fresh #2) to the 7-days frozen SynCom (Table 3).

**Table 3.** Adjusted p-values of ANOVA followed by Tukey HSD tests comparing CFUs/g fresh weight on maize roots collected 10 days post-emergence for each SynCom species between two fresh, and frozen (1 h, and 7 d) replicated experiments (data are in Table 1). Significantly different comparisons (p<0.05) are marked with an asterisk (*).

|  |  | SynCom Member |  |  |  |  |  |  |
| --- | --- | --- | --- | --- | --- | --- | --- | --- |
|  |  | *S. maltophilia* | *B. pituitosa* | *C. pusillum* | *E. ludwigii* | *C. indologenes* | *H. robiniae* | *P. putida* |
| Experiment #1 | Fresh #1 v 1h | 0.56 | 0.99 | 0.49 | 0.12 | 0.15 | 0.95 | 0.12 |
|  | Fresh #1 v 7d | 0.47 | 0.99 | 0.01* | 0.57 | 0.99 | 0.40 | 0.41 |
| Experiment #2 | Fresh #2 v 1h | 0.84 | 0.50 | 0.64 | 0.03* | 0.90 | 0.32 | 0.87 |
|  | Fresh #2 v 7d | 0.002* | 0.000001* | <1 e-7* | 0.00002* | 0.04* | 0.0002* | 0.003* |
|  | Fresh #3 v 1h | 0.21 | 0.0003* | <1 e-7* | 0.08 | 0.001* | 0.38 | 0.16 |
|  | Fresh #3 v 7d | 0.56 | 0.95 | 0.94 | 0.95 | 0.96 | 0.70 | 0.75 |

To test the impact of longer-term freezing on species culturability in the SynCom master mix, we plated read-to-use SynCom stocks on selective media after 60 days of storage at -80°C. The expected titer of each species in the frozen SynCom after dilution with 8.3 ml of PBS was 10^6^ for *S. maltophilia* and *P. putida* and 10^7^ CFU/ml for the other five species. Both *S. maltophilia* and *H. robinae* showed much lower CFU numbers than expected (1-2 orders of magnitude lower) (Figure 2).

**Figure 2.**
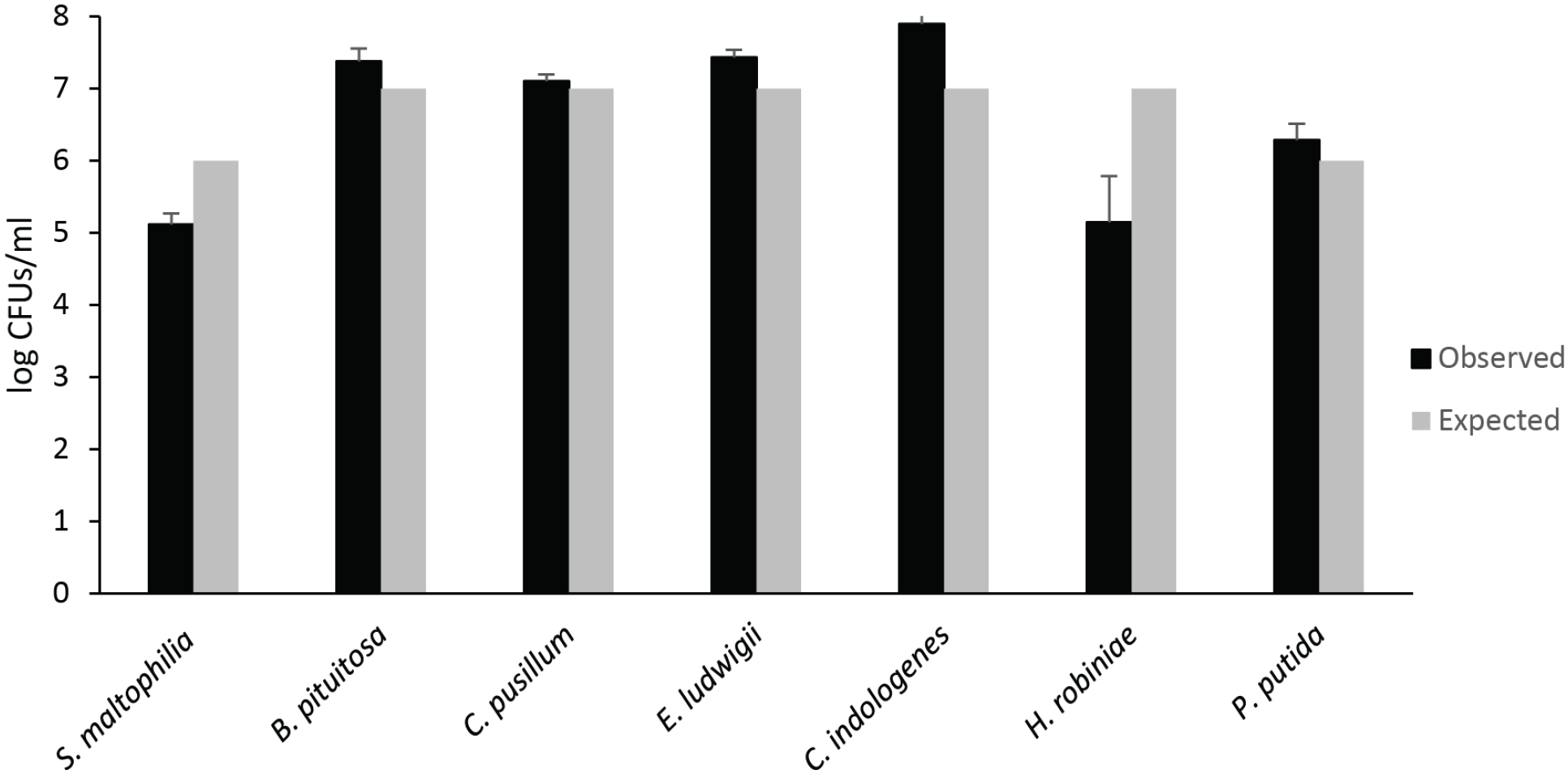
Recovery of species from frozen SynComs by plating on selective media following storage at -80°C for 60 days.

### Relative abundance of SynCom members

Despite variation in the absolute abundance of each community member (CFU/g of fresh root) using both frozen and fresh SynComs for seed inoculation (Tables 1-3), the relative abundance of each SynCom member colonizing the root remained consistent across all of the SynComs whether fresh or frozen (Figure 3). The exception was *C. pusillum*, which showed significant differences (ANOVA followed by Tukey HSD, p<0.05) between different mixes; however, these differences were not associated with a specific SynCom treatment. In other words, lower relative abundances for *C. pusillum* were observed in both fresh and frozen SynComs.

**Figure 3.**
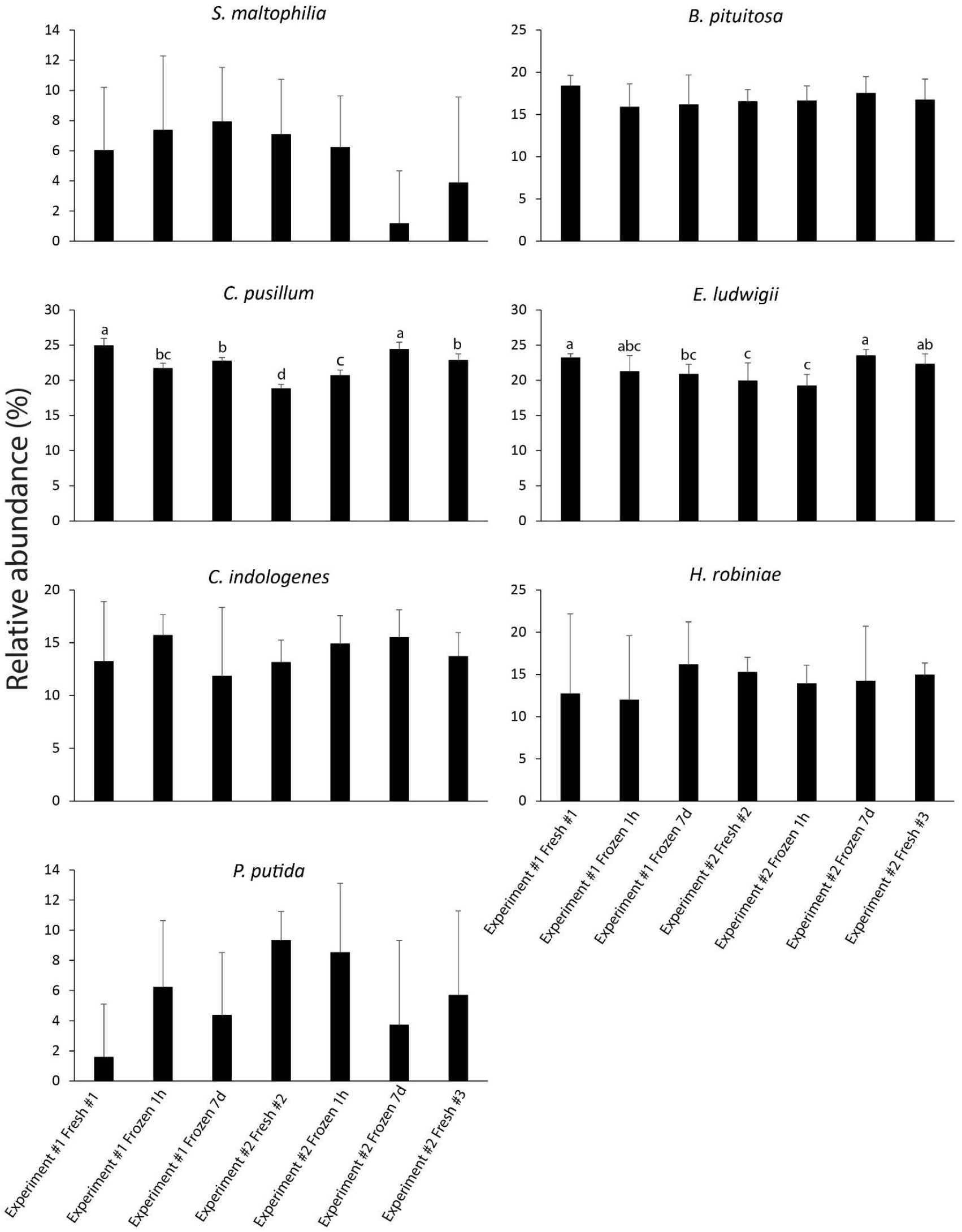
Percent relative abundance of each member of the SynCom based on CFUs/g of fresh corn root 10 days post emergence. Bars with the same letter above the bar were not significantly different (p < 0.05). One-way ANOVA followed by Tukey HSD was performed for each species, p-values were adjusted for multiple comparisons using the Benjamin Hochberg method. Error bars represent standard deviations. Bars without letters above were not significantly different for any of the comparisons within the same species (p > 0.05).

## Discussion

Our goal with this study was to evaluate the potential of freezing aliquots of SynComs in lieu of constructing a new community for each experiment. Freezing aliquots of a SynCom master mix provides two benefits to SynCom-host studies. First, it reduces the amount of work required to culture individual members in order to construct the community each time an experiment is initiated, and second, our results suggest that consistency of species relative abundances in frozen aliquots is equal to if not better than the consistency of re-culturing and constructing a fresh community for each experiment.

Although the same protocol was used to prepare each replicated SynCom, the absolute abundances of members of each SynCom harvested from maize roots significantly differed in each of the three communities. The variation in each member of the freshly-constructed SynComs harvested from maize roots was quite high, ranging from standard deviation log-values of 0.18 (*E. ludwigii* and *C. pusillum*) to 2.93 (*H. robiniae*) (Table 1). Several SynCom studies report the log-scale variance in colonies on the root surface across replicates. For example, the original study of the SynCom tested here reports the range of variance for colonization of each of the seven community members (Niu et al., 2017). Niu et al. reported that replicates collected at the same time point had a +/-log variance anywhere from 0.04 (SD for *E. ludwigii*) to 1.78 (SD for *P. putida*) (Niu et al., 2017). These values are consistent with another study of a 12-member SynCom in maize that indicates colonization variance between a +/-variance of 0.5-1.0 log (Figueiredo dos Santos et al., 2021). Another study that explored cryo-preservation of a 17-member SynCom also showed large differences in community composition and root colonization in *Brachypodium* (Coker et al., 2022). Despite the high variance that we have presented here, and which was reported in other studies, the variation in colonization is not reported in most published SynCom studies even though it is potentially important in understanding microbial colonization of plants.

We think that part of the significant differences observed in absolute species abundances are likely due to variation introduced during harvesting and plating for counting, rather than variability due to community construction or freezing. In Experiment 1 all the root samples (fresh, frozen 1h and 7d) were processed at the same time. In this Experiment, only one out of 14 comparisons between frozen SynCom numbers and fresh SynCom numbers was significant. In Experiment 2, root samples inoculated with fresh mix #3 and 7-day frozen mix were harvested and processed at the same time and none of the comparisons was significant. While all comparisons of the fresh mix #2, which was harvested earlier, to the 7-day frozen mix were significant (Table 3).

Despite the high variability in absolute abundance of individual community members harvested from maize roots across each SynCom, the relative abundances of SynCom members were similar to each other and to the relative abundances reported previously (Niu et al., 2017). This indicates that potential variation in microbial species ratios introduced by SynCom treatment did not impact overall colonization patterns of SynCom members on corn roots. However, since we did not measure ratios of SynCom members for each treatment by plating inocula prior to application to corn seeds, we do not know how much variation was present in SynCom member ratios between treatments. The observed stability in relative ratios after colonization could thus either be due to limited variation of species ratios in the inoculum or deterministic establishment of specific relative ratios upon root colonization driven by interactions with the plant and/or other SynCom members independent of input ratios. The literature on inoculation of plant and animal hosts with SynComs supports both scenarios. For example, Carlström et al. (2019) showed that ratios of 62 SynCom members in the inoculum were not predictive of ultimate colonization patterns in the *Arabidopsis* phyllosphere and some SynCom members consistently colonized to high relative abundances, while others consistently failed to colonize. The data from Carlström et al. indicates that at least in part colonization patterns are deterministic and independent of inoculum ratios. On the other hand, Venturelli et al., (2018) showed that ratios of SynCom members in the inoculum can impact the final community in a human gut microbiome SynCom. They demonstrated that after 72 h of cultivation, 12% of the communities displayed legacy dependence on the initial ratio (Venturelli et al., 2018). In summary, while in our study the relative colonization ratios of SynCom members on corn roots were similar across all treatments, it will be important to assess the impact of strongly shifted SynCom member ratios in the inoculum on root colonization patterns as such ratio shifts might be caused by longer-term storage of ready-to-use SynCom stocks.

Our study has at least two limitations, which should be addressed in future work. First, we did not investigate how long ready-to-use stocks can be stored in the freezer before losing their ability to colonize corn roots at reproducible relative abundances. Ideally, ready to use inocula allow for reproducible inoculation for at least 1 year or more to enable execution of multiple repeat experiments and potentially sharing of ready-to-use stocks with other members of the scientific community for reproducibility across laboratories. Second, often experiments with SynComs require constructing SynComs with different compositions such as for example the dropping out of suspected keystone species. For these types of experiments it still would be beneficial to not have to grow all the community members *de novo* every time. Thus mixing of SynComs from ready-to-use stocks of individual SynCom members or groups of SynCom members would be ideal. However, this has to our knowledge not been tested yet in any system and thus would require careful testing and validation.

Finally, although we only studied the applicability of a frozen ready-to-use SynCom for this particular maize SynCom (Niu et al., 2017), we have demonstrated the utility of frozen SynCom stocks in terms of reducing labor and improving inoculum consistency between experiments. The principle works and the approach used here can be applied to validate frozen ready-to-use mixes for other SynComs.

## Author contributions

All authors contributed to the conception and design of this study. SV and CT ran experiments. SV and JJP analyzed the data. JJP, SV and MK wrote the first draft of the manuscript. All authors contributed to manuscript revision, read, and approved the submitted manuscript.

## Funding

This work was supported by the United States National Science Foundation under award number IOS-2033621 (M. Wagner and M. Kleiner), the U.S. Department of Agriculture National Institute of Food and Agriculture under award number 2022-67013-36672 (M. Kleiner and M. Wagner) and the Novo Nordisk Foundation INTERACT project under award number NNF19SA0059360 (M. Kleiner).

## Acknowledgements

We are grateful to Dr. Heather Maughan for feedback on the manuscript.

